# Arginine-rich C9ORF72 ALS Proteins Stall Ribosomes in a Manner Distinct From a Canonical Ribosome-Associated Quality Control Substrate

**DOI:** 10.1101/2022.02.09.479805

**Authors:** Viacheslav Kriachkov, Hamish E G McWilliam, Justine D Mintern, Shanika L Amarasinghe, Matt Ritchie, Luc Furic, Danny M Hatters

**Affiliations:** Department of Biochemistry and Pharmacology and Bio21 Molecular Science and Biotechnology Institute. The University of Melbourne, Victoria 3010 Australia; Department of Microbiology and Immunology, Peter Doherty Institute of Infection and Immunity, The University of Melbourne, Parkville, Victoria, Australia; Walter and Eliza Hall Institute of Medical Research, Victoria 3052 Australia; Translational Prostate Cancer Research Laboratory, Peter MacCallum Cancer Centre, Melbourne, Victoria, Australia, 3000; Sir Peter MacCallum Department of Oncology, University of Melbourne, Parkville, Victoria, Australia 3010; Monash Biomedicine Discovery Institute Cancer Program, Department of Anatomy and Developmental Biology, Monash University, Clayton, Victoria, Australia, 3800

## Abstract

Hexanucleotide expansion mutations in *C9ORF72* are a cause of familial amyotrophic lateral sclerosis. We previously reported that long arginine-rich dipeptide repeats (DPR), mimicking abnormal proteins expressed from the hexanucleotide expansion, caused translation stalling when expressed in cell culture models. Whether this stalling provides a mechanism of pathogenicity remains to be determined. Here we explored the molecular features of DPR-induced stalling and examined whether known regulatory mechanisms of ribosome quality control (RQC) are involved to sense and resolve the stalls. We demonstrate that arginine-containing DPRs lead to stalling in a length dependent manner, with lengths longer than 40 repeats invoking severe translation arrest. Mutational screening of 40×Gly-Xxx DPRs shows that stalling is most pronounced where Xxx are positively charged amino acids (Arg or Lys). Through a genome-wide knockout screen we find that genes regulating stalling on polyadenosine mRNA coding for poly-Lys, a canonical RQC substrate, respond differently to the readthrough of arginine-rich DPRs. Indeed, we find evidence that DPR-mediated stalling has no natural regulatory responses even though the stalls may be sensed, as evidenced by an upregulation of RQC gene expression. These findings therefore implicate arginine-rich DPR-mediated stalled ribosomes as posing a particular danger to cellular health and viability.

## INTRODUCTION

GGGGCC hexanucleotide repeat expansion mutations in intron 1 of *C9ORF72* are the cause of familial amyotrophic lateral sclerosis (ALS) and frontotemporal dementia (1,2). Normally, there are less than 24 repeats, whereas the length is expanded to often hundreds in ALS-causing alleles (3). The expanded GGGGCC leads to the production of sense and antisense mRNA products that display two unusual features that have been postulated to contribute to disease. One is that the mRNA can form granular intracellular foci that contribute to toxicity through RNA-based mechanisms (4). The other is that both sense and antisense mRNA can be translated through alternative initiation codons (i. e. non-AUG translation) to produce five distinct dipeptide-containing polymers (poly-GA, GR, GP, PR and PA) (5). These abnormal proteins accumulate in the brain of patients with *C9ORF72* mutations. The two arginine containing dipeptide repeats (DPR), poly-GR and poly-PR, have been shown to be particularly toxic when added to cells or when expressed in cellular and organismal models (6–8). The toxicity is preserved if the GR and PR are encoded by mixed codons, suggesting that the protein sequence itself is directly toxic, and therefore not entirely arising from the mRNA (7,8).

The fidelity of protein synthesis involves mechanisms that detect and eliminate spontaneous cases of aberrant translation (9). This includes: translation of mRNAs with defects including mRNAs with stable stem-loop structures, damaged bases or other obstacles to elongation (no-go decay, NGD); mRNAs that lack stop codons (nonstop decay, NSD) and mRNAs with premature stop codons (nonsense-mediated mRNA decay, NMD) (10). A marked decrease in elongation rates on such mRNAs can lead to formation of persistent ribosomal collisions, which are targeted for ribosome-associated quality control (RQC) pathway that resolves such stalls and causes degradation of both defective mRNA and polypeptide (11–14). In the event stalls happen aberrantly and remain unresolved they can have pathological consequences. It was reported that simultaneous mutation in CNS (central nervous system)-specific tRNA gene that leads to stalling on specific codons and loss of GTPBP2, a ribosome rescue factor resolving such stalls, causes widespread neurodegeneration in mice (15). Similarly, mice with a loss of function of LTN1 protein that targets stalled nascent chains for degradation also exhibited a neurodegenerative phenotype (16).

Recent studies from us and others showed that expression of mRNA encoding long repeats of poly-GR and poly-PR causes severe ribosomal stalling (17,18) and it was proposed that the positively charged R-rich nascent polypeptide electrostatically jams the ribosome exit tunnel (19). However, the mechanisms by which stalling occurs and whether it contributes to toxicity remain unclear. Another outstanding question is whether known mechanisms that target stalled ribosomes in cells are able to sense and modify stalls caused by poly-GR and poly-PR. Here we examine the molecular features of stalling from poly-PR. We examined the length of the repeat required to stall and specificity of arginine in a context of DPR-mediated arrest as well as conducted a screen for potential genetic regulators of stalling on poly-PR protein. Collectively our findings indicated that the mechanisms involved in stalling caused by long R-rich DPRs are unequivocally distinct to the mechanisms leading to stalling on poly-K coded by polyadenosine sequence that mimics defective mRNAs lacking stop codons. Our findings point to a mechanism of toxicity that relates to a lack of an evolved capacity to resolve stalls involving R-rich DPRs.

## MATERIALS AND METHODS

### DNA constructs

Plasmids coding linker, poly-K, 102×GR and 102×PR in dual fluorescence stall reporter were prepared as described (17). cDNA for 20×, 30×, 40×, 50×, 75× GR and PR, and 40× GA, GC, GD, GE, GF, GH, GI, GK, GL, GM, GN, GP, GQ, GS, GT, GV, GW, GY were synthesized and cloned into dual fluorescence stall reporter (GenScript). For knockout cell pools, sgRNA sequences targeting genes of interest were designed using Benchling software. They were cloned into lentiCRISPRv2 vector (a gift from Feng Zhang; Addgene plasmid # 52961) (20) using BsmBI restriction sites, as outlined in the protocol from Zhang lab (available at Addgene). For stable Tet-inducible cell lines, GFP-P2A-ChFP, GFP-P2A-poly-K and GFP-P2A-102×PR were cloned by restriction digestion and ligation and inserted into pLVX-TetOne-Puro vector (Takara Bio # 631849) using Gibson assembly. All cloned constructs were validated by sequencing. DNA preparations were made with Stbl3 *E. coli* cells (Thermo Fisher). The sequence information is available (Table S1).

### Cell Culture

HEK293T cells, obtained originally from the American Type Culture Collection (ATCC, Manassas, Virginia), were maintained in complete DMEM (Dulbecco’s modified Eagle medium (DMEM) supplemented with 10% v/v fetal calf serum and 1% GlutaMAX). Cells were cultured in a humidified incubator with 5% v/v atmospheric CO^2^ at 37 °C. For the drug treatment experiments, harringtonine (Abcam # ab141941; 20 mM working stock in DMSO) or cycloheximide (Sigma-Aldrich # 01810-1G; 50 mM working stock in DMSO) were added to cells 6 hours post-transfection alongside a change of culture medium, and cells were analyzed by FACS at 24 hours post-transfection.

### Ribosomal stalling assays

HEK293T cells were transfected with staller constructs using Lipofectamine 3000 reagent and harvested 24 h post-transfection. Cells were analyzed using LSRFortessa X-20 flow cytometer (BD Biosciences). Side and forward scatter height, width, and area were collected to gate for single live cell population. GFP fluorescence were collected with the 488-nm laser and FITC (530/30) filter to gate for transfected cells, ChFP fluorescence was collected with the 561-nm laser and mCherry (610/20) filter. Flow cytometric gating and data analysis was performed using FlowJo software (v10.5.3) and graphs were analyzed in RStudio (v. 1.4.1106) and GraphPad Prism 8. Median FITC-A and median mCherry-A fluorescence values were used to calculate ChFP/GFP ratio for each sample. All ratios were normalized to average of ChFP/GFP ratios for fluorescence reporter with linker sequence (negative no-stall control).

### Confocal imaging

HEK293T cells were transfected with GFP-tagged poly-PR constructs using Lipofectamine 3000 reagent. 24 hours after transfection, cells were fixed in 4% paraformaldehyde for 15 min at room temperature. Nuclei were counterstained with Hoechst 33342 at 1:750 dilution (Thermo Fisher # H3570) for 20 min then washed twice in PBS. Confocal images were obtained on Zeiss Elyra LSM880 confocal microscope using 63× oil-immersion objective lens at room temperature. Single color controls were used to establish and adjust to remove bleed through of the emission filter bandwidths. FIJI version of ImageJ (v.1.53c) (21) was used for image processing.

### Lentiviral work

Lentivirus for Brunello knock out library, single knock out cell lines and Tet-inducible cell lines were made per Thermo Fisher guidelines with minor modifications. HEK293T cells were transfected with pCMV-VSV-G (a gift from Bob Weinberg; Addgene plasmid # 8454) (22), psPAX2 (a gift from Didier Trono; Addgene plasmid # 12260) and lentiviral transfer plasmid in equimolar ratios using Lipofectamine 3000 reagent. Virus-containing media was collected 48 hours post-transfection and filtered using 0.45 μm filter. Virus supernatant was stored at –80°C. HEK293T cells were transduced with virus in the presence of 8 μg/mL polybrene. The media was refreshed with complete DMEM 24 h after transduction. At 48 h after transduction, antibiotic selection for infected cells were performed with 1 μg/ml puromycin (Thermo Fisher # A1113803; 10 mg/mL working stock in 20 mM HEPES buffer) for 3 days.

### RNA sequencing

HEK293K cell lines harbouring the constructs under the Tet-on expression system were induced for expression by adding complete DMEM media containing 500 ng/μl doxycycline (Sigma-Aldrich # D9891; 1 mg/ml working stock in DMSO) to the cells for 48 h prior to harvesting. Expression was confirmed by microscopy and flow cytometry for the fluorescent proteins. For each construct, non-induced cells were collected in parallel to the doxycycline-induced cells. Cell were collected in ice-cold lysis buffer (20 mM Tris pH 7.4, 150 mM NaCl, 5 mM MgCl^2^, 1 mM DTT, 100 μg/ml cycloheximide, 1% v/v Triton X-100 and 25 U/ml Turbo DNase I), triturated ten times through a 26-gauge needle and centrifuged for 10 min at 20,000 *g* at 4 °C. The RNA concentration in the supernatant was measured using Promega QuantiFluor RNA System. Total RNA was isolated by Zymo Research Direct-zol RNA MiniPrep Kit. The stranded mRNA library was prepared and sequenced on Illumina Novaseq 6000 100bp SR by the Australian Genome Research Facility. Triplicate samples were prepared. Count tables were generated in Galaxy.org using HISAT2 (v.2.1.0) algorithm (23), and the list of differentially expressed genes were generated with DESeq2 package (v.1.30.1) (24) in R (v.4.0.5). Gene set enrichment analysis (GSEA) was performed using a javaGSEA desktop application (v.4.1.0) (25,26). Curated gene set GSEA: C5.GO.BP.v7.4 was obtained from the Molecular Signatures Database (MSigDB) (27). Data sets with FDR < 0.1 were considered as statistically significant. Enrichment map visualization was performed in Cytoscape (v.3.8.0) (28).

### Genome-wide CRISPR knockout screening

The human CRISPR knockout pooled library Brunello was obtained from Addgene (a gift from David Root and John Doench; Addgene # 73178) (29). The experiment was done as outlined in the following protocol (30), with minor modifications. For each of 3 replicates, HEK293T cells were transduced at MOI = 0.4. Following antibiotic selection, at least 10^8^ cells were transfected with either poly-K or 102×PR staller. 24 hours post-transfection half of cells were subjected to sorting while the other half was left as unsorted control population. Cells were sorted at BD InFlux cell sorter for top 5% of high and low ChFP/GFP ratio populations. gDNA from collected cells was extracted with salt precipitation protocol described here (31). The sgRNA-containing cassettes were amplified as described previously (32). The pooled amplicons were sequenced using an Illumina NovaSeq 6000. The results were analyzed and false discovery rate (FDR) and log2 fold change (LFC) values for each gene were calculated using MAGeCK (v.0.5.9.2) computational tool (33). Functional enrichment analysis was done in Cytoscape (v.3.8.0) for all screen hits with FDR<0.05.

### Generation of knockout cell pools

Single sgRNA expression vectors were cloned and lentivirus production with the following transduction was performed as described above. More than 50% knockout efficiency for each pool was validated by ICE-seq analysis (34) after isolating total gDNA and sequencing the targeted region of gene of interest. The sgRNA sequences targeting each selected gene used are shown in Table S1. Cells expressing a non-targeting sgRNA were used as a control pool. Fold changes in ChFP/GFP ratio for poly-K, 102×PR or 102×GR staller after individual gene knockout were calculated by dividing the ratio from cells expressing sgRNA to ratio from cells without sgRNA. For each knockout pool, fold changes in ChFP/GFP on a staller sequence were compared against fold changes in ChFP/GFP on a linker sequence that does not cause stalling.

### Statistical analysis

Details of quantification including number of samples analyzed and statistical tests used are described in the corresponding figure legends and methods section.

## RESULTS

### Arginine-rich dipeptide repeats stall during translation in a length-dependent manner

To find the dipeptide repeat length at which the nascent chain is long enough to hinder ribosome readthrough, we used a series of DNA sequences coding PR and GR dipeptide repeats of various lengths: 10×, 20×, 30×, 40×, 50×, 75×, and 102×. All DPR-coding sequences were codon-optimized to minimize repetitive mRNA sequences and codon repetitiveness. Readthrough efficiency was assessed using the previously described dual-reporter transcript (Figure 1A) (35–38). Briefly, this reporter consists of mRNA sequence to be tested for stalling which is flanked, in frame and without stop codons, with sequences encoding green fluorescent protein (GFP) and mCherry fluorescent protein (ChFP). P2A sequences separate the test sequence from the GFP and ChFP and cause the ribosome to “skip” the glycyl-prolyl peptide bond formation. As such, a single mRNA transcript can translate three separate proteins (39). The yield of ChFP versus GFP allows a measure of whether the test sequence can modulate stalling, i.e. the ChFP/GFP ratio will be reduced in the event stalling occurs. While both poly-PR and poly-GR proteins lowered the ChFP/GFP ratio progressively in a length-dependent manner suggesting increased levels of stalled translation, they also showed different trends, with poly-GR appeared to stall more substantially at the longer lengths of dipeptide repeat (Figure 1B, C).

**Figure 1.**
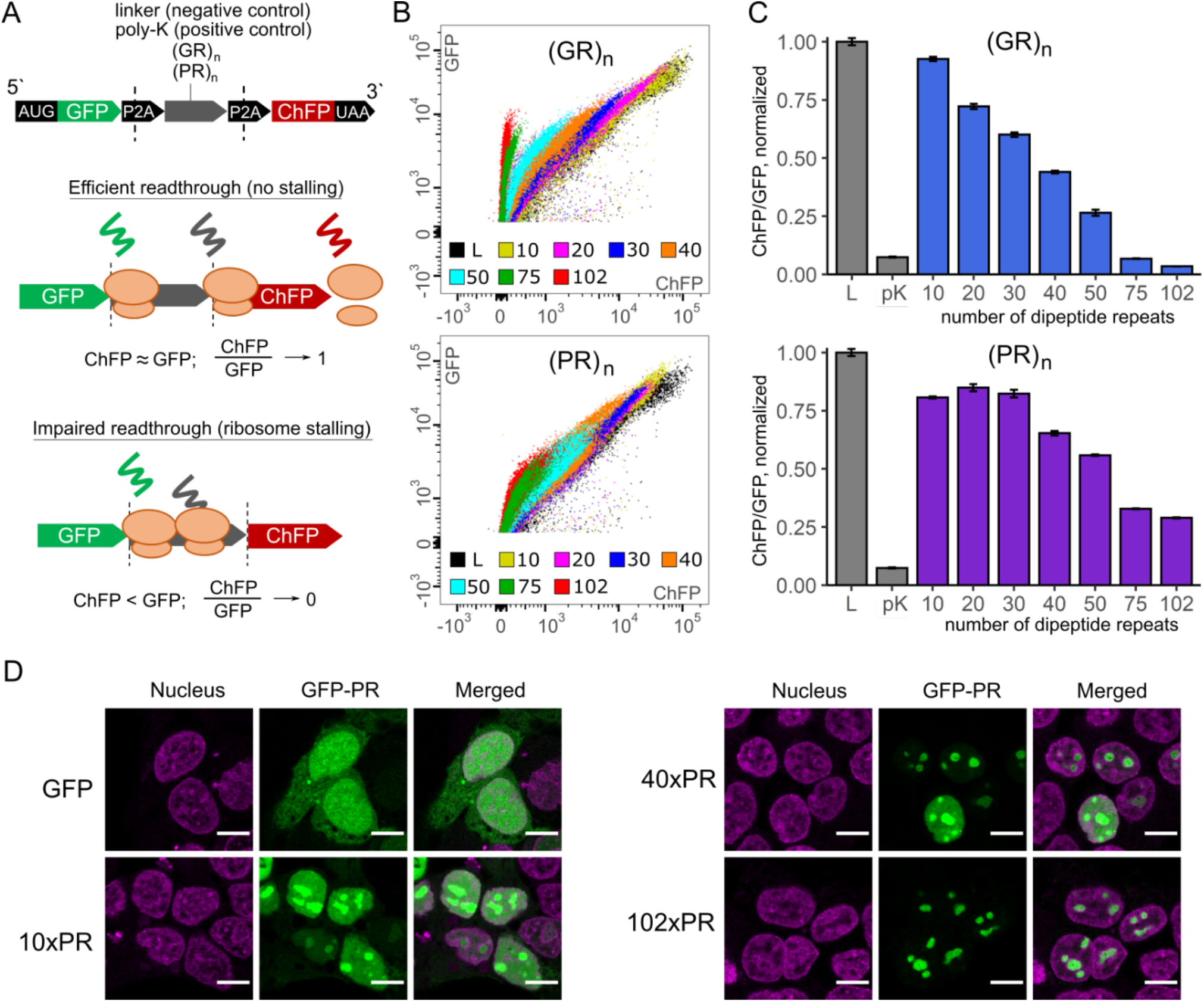
Ribosome stalling assays with poly-GR and poly-PR of different lengths. (**A**) A schematic of the sequence of the dual-fluorescent stall reporter. (**B**) Flow cytograms of HEK293T cells after 24 h of transfection with poly-GR and poly-PR coding sequences of indicated length (10×, 20×, 30×, 40×, 50×, 75× or 102×) inserted into fluorescent reporter, L – linker. (**C**) ChFP/GFP ratios for different poly-PR and poly-GR stall reporters. L – linker, pK – poly-K coded by (AAA) codons. Median GFP and ChFP fluorescence values were measured by FACS and resulted ChFP/GFP ratios were normalized to ChFP/GFP ratio of negative stall reporter (linker). Bars show means and SD, n=3. (**D**) Confocal images of HEK293T cells after 24 h transfection with GFP or GFP tagged with 10×PR, 40×PR and 102×PR. The nucleus was stained with Hoechst 33258. Scale bar represents 10 μm.

Poly-PR is known for its property to form nuclear foci (7) and to test if this phenotype changes with polymer length, we expressed 10×PR, 40×PR and 102×PR fused to C-end of GFP in HEK293 cells (Figure 1D). Notably, non-staller GFP-10×PR, weak GFP-40×PR staller and stalling-induced GFP-102×PR had similar localization patterns. Overall, this finding suggests that poly-PR confers a functional property for recruitment to bodies in the nucleus, however, this feature appears independent to stalling or to the repeat length.

### Specificity of arginine in the context of DPR-mediated ribosome stalling

Prior work has suggested that positively charged amino acid patterning can influence the stalling of translation during synthesis (40). Given that other non-arginine DPRs associated with *C9ORF72* hexanucleotide expansion (poly-PA and poly-GA) do not appear to lead to stalling (17), we next examined if stalling in a DPR context was specific to arginine. A small library of DPRs was assessed in the stalling assay whereby each variant was a DPR containing one of the other 19 canonical amino acids alternating with glycine. The sequences were codon optimized to minimize secondary structures and codon usage bias where possible. The DPRs containing charged residues at neutral pH (Arg, Asp, Glu and Lys) led to the largest decreases in ChFP/GFP ratios, suggestive of substantial ribosome stalling (Figure 2A–B, Figure S1). Of these, the negatively charged DPRs (40×GD and 40×GE) appeared more severe in their effect than the positively charged DPRs (40×GR and 40×GK).

**Figure 2.**
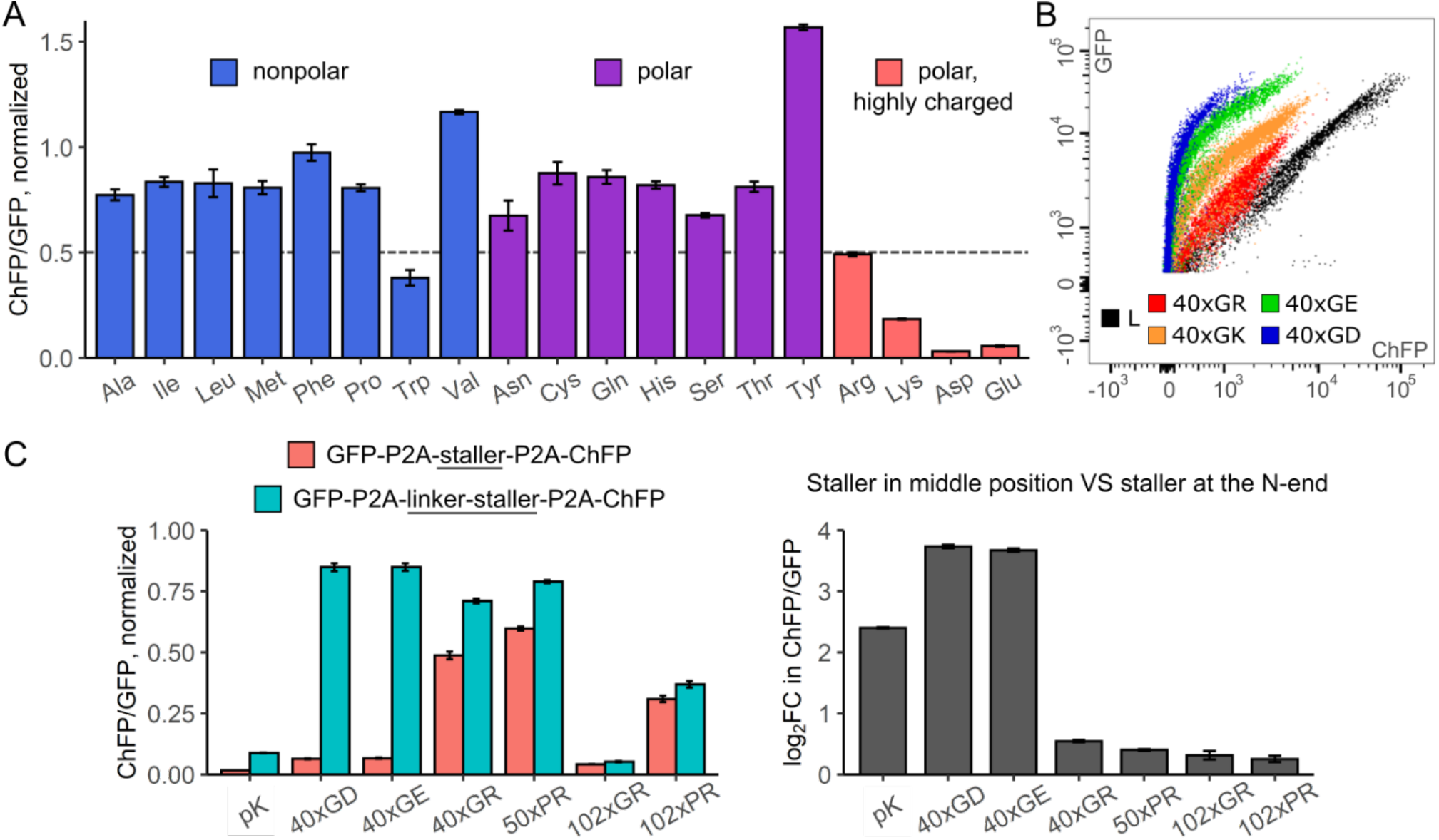
Ribosome stalling assays with library of different dipeptide repeats. (**A**) The relative ChFP/GFP ratios for different DPRs normalized to ratio in cell transfected with negative staller control (linker). Each DPR is 40×Gly-Xxx, where Xxx – amino acid residue displayed on x axis. Bars show means and SD, n=4. (**B**) Flow chart showing decrease in ChFP readout from all fluorescent reporters containing charged DPRs: 40×Gly-Arg (40×GR), 40×Gly-Lys (40×GK), 40×Gly-Glu (40×GE), 40×Gly-Asp (40×GE). L – linker. (**C**) Left plot: the relative ChFP/GFP ratios (normalized to linker) for different DPR stallers and poly-K (pK) that either start right after P2A sequence (pink bars) or have a linker sequence added to their N-end (blue bars). Bars show means and SD, n=3. Right plot: log2 fold changes in ChFP/GFP ratio after adding the linker sequence between P2A and N-end of DPR.

It was previously found that the translation of open reading frames containing multiple acidic residues at the start of the sequence results in intrinsic ribosomal destabilization, which leads to translation abortion in *E. coli* (41). Such a mechanism, which is distinct to stalling, may lead to selective reduction of ChFP in our assay and therefore confound interpretation of stalling. To examine for this possibility, we inserted a linker sequence at the N-terminus of the DPR sequences of acidic DPRs and the R-rich DPRs. Both the 40×GD and 40×GE constructs no longer appeared to stall under these conditions, whereas the stalling was preserved for 102×PR and 102×GR constructs (Figure 2C). We therefore concluded that stalling in the context of DPR is selective to positively charged (arginine and lysine) moieties.

### poly-PR expression leads to significant changes in the expression of various stress response genes including the upregulation of RQC genes at transcriptional level

To determine whether polyadenosine-mediated stalling induces similar change in gene expression as poly-PR stalling, we performed transcriptome analysis. We created three stable HEK293T cell lines expressing GFP-P2A-ChFP (which provides a negative control with no stalling), GFP-P2A-poly-K, whereby poly-K was coded with (AAA) codons (and therefore represents the canonical RQC substrate, which induces stalling), and GFP-P2A-102×PR under a Tet-inducible promoter (Figure 3A). Gene expression levels were compared for the poly-K or 102×PR staller versus the ChFP negative control. Principle component analysis indicated that 102×PR-expressing cells clustered separately to all the other sample groups as well as the cells that were uninduced (Figure 3B). Volcano plots revealed that expression of polyadenosine led to relatively minor changes in gene expression, whereas 102×PR expression led to substantial changes (Figure 3C). Indeed, 102×PR expression resulted in 9962 genes significantly changing expression compared to cells expressing ChFP. Of these, 5086 genes were significantly upregulated and of these 1037 more than 2-fold. 4876 genes were significantly down-regulated and of these 647 genes more than 2-fold (Table S2). For the poly-K staller, 303 genes were significantly upregulated, but none more than 2-fold. 102 genes were down-regulated, and of these only 2 showed more than 2-fold decrease.

**Figure 3.**
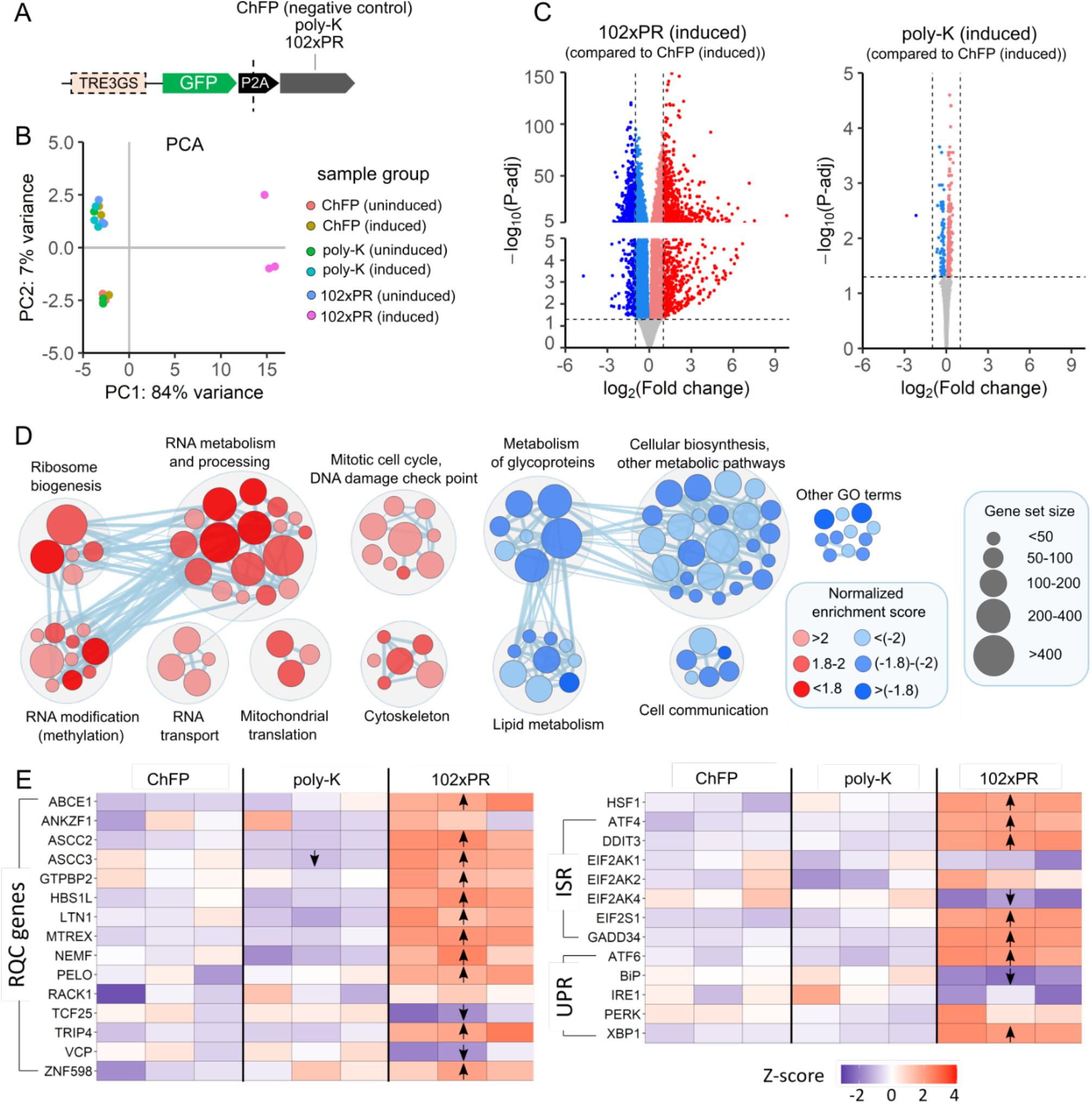
Changes in global transcriptome caused by expression of poly-K or poly-PR stallers. (**A**) A schematic of the transcript used to generate stable cell lines. TRE3GS – promoter containing tetracycline response element (TRE). (**B**) Principal component analysis plot for all samples. Each group was done in triplicate (n = 3). (**C**) Volcano plots showing differentially expressed genes (p-adj < 0.05, horizontal line) for each staller. Red – upregulated genes, blue – downregulated genes. Vertical lines represent cut-off values for log2(Fold change): 1 and −1. (**D**) Enrichment map with GO terms that were significantly up-(red) or downregulated (blue) upon 102xPR expression (FDR q-value < 0.1) and grouped into the following clusters. Color intensity is proportional to the normalized enrichment score (NES) for a given GO term. Node size is proportional to a total number of genes in GO category (gene set size). (**E**) Heat maps with Z-scores for selected genes from various stress pathways (RQC – ribosome quality control, ISR – integrated stress response, UPR– unfolded protein response) in cells with induced expression of ChFP, poly-K coded by (AAA) codons or 102×PR. Arrows indicate differentially expressed genes (up/down arrows for up- and down-regulated genes respectively) in cells with induced staller expression when compared to ChFP-expressing cells.

Gene set enrichment analysis identified 170 Gene Ontology (GO) terms that were significantly upregulated in the 102×PR-expressing cells and 327 GO terms that were downregulated (Table S2). Further pathway enrichment analysis (Figure S2) revealed that many GO terms related to ribosome biogenesis, RNA metabolism, RNA processing, RNA transport and RNA modification (mostly methylation) were upregulated in response to 102×PR expression (Figure 3D). It has been reported that the expression of poly-PR inhibits global translation possibly due to interactions of poly-PR with ribosomes and other components of the translation machinery (42–44). As such, our results may reflect a compensatory response to activate genes involved in global translation. Other upregulated GO terms were linked to cytoskeleton which is in the agreement with prior findings of poly-PR disturbing the organization of the cytoskeleton (45) and interacting with actin-related cytoskeletal proteins (17). Our data also supports the role for poly-PR in inhibiting DNA double strand break repair pathways (46) as we saw an upregulation of GO terms related to mitotic cell cycle such as negative regulation of nuclear division (GO:0051784), negative regulation of metaphase/anaphase transition of cell cycle (GO:1902100), negative regulation of chromosome organization (GO:2001251) and mitotic intra-S DNA damage checkpoint signaling (GO:0031573).

In addition, many metabolic pathways were downregulated as a result of 102×PR expression. Prominent were clusters of GO terms related to glycoprotein metabolism, lipid biosynthesis and other terms related to cellular metabolic processes. These findings suggest substantial metabolic adaptation to the 102×PR expression. This result is consistent with previous reports finding that poly-PR directs promiscuous proteome binding and cytotoxocity (17,43). The other point of note was the significant upregulation of RQC-associated genes as well as some genes involved in integrated stress response (ISR) (Figure 3E). These effects were not seen with the poly-K construct. This finding suggested that stalling of poly-K coded by polyadenosine can be readily accommodated by proteins that already present in cell whereas 102×PR stalled complexes are more detrimental to cell health and require a massive stress response from a cell.

### Genome-wide knockout screen unmasks a distinct signature of regulators of 102×PR readthrough compared to poly-K readthrough

To investigate whether mechanisms that are known to regulate stalls on polyadenosine mRNA are involved in sensing and responding to poly-PR stalls, we performed a genome-wide CRISPR/Cas9 knockout screen using the Brunello library, which targets 19,114 human genes at a coverage of 4 gRNA targets per gene (29). We performed two screens: one with the 102×PR staller reporter and the other with poly-K coded by (AAA) codons as a reference for known RQC regulatory mechanisms (37,47,48). Flow cytometry was used to sort and collect two groups of cells reflecting the higher and lower 5% ranges of ChFP/GFP ratios (Figure 4A). Population “H” represented cells with the highest ChFP/GFP ratios, which should encompass the highest readthrough levels. Population “L” represented cells with lowest ChFP/GFP ratio, which should encompass the highest levels of stalling. The logic of the screen was that a knock-out of a particular gene involved in the resolution of stalling or RQC function would increase stalling (i.e. be over-represented in the L population). Conversely, gene knockouts of genes involved in pausing translation on stalled sequences, or recruitment of machinery to stalls, would be expected to be enriched in the H population, which depicts higher levels of readthrough.

**Figure 4.**
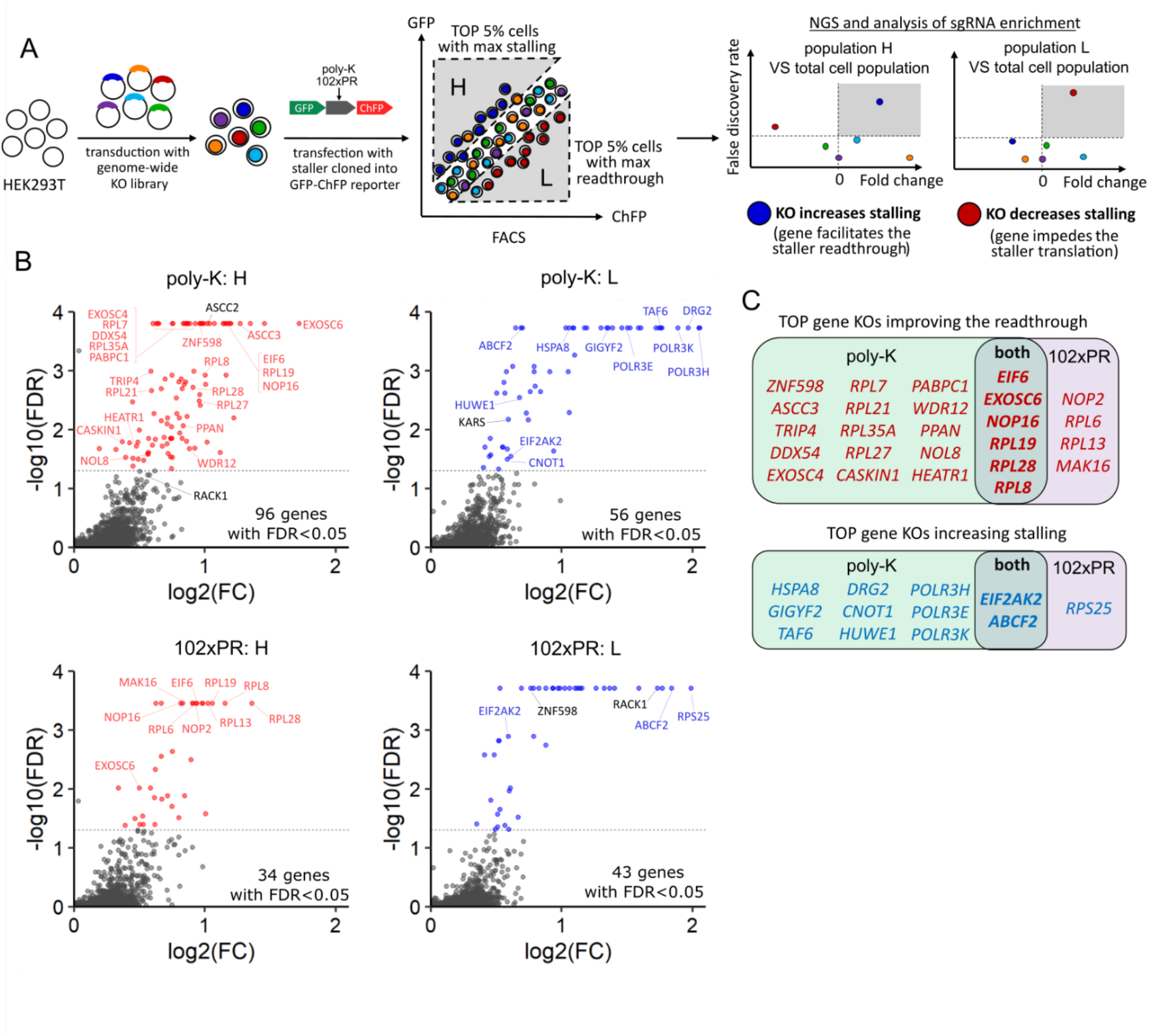
Genome wide knockout screening for genes modulating stalling of poly-K and poly-PR stallers. (**A**) Schematic of strategy. (**B**) Volcano plots of gene KO enrichment. The number on the plot indicates the number of screen hits (FDR < 0.05) for each category. Genes which targeting sgRNAs were enriched in population H (highest ChFP/GFP ratio) are shown in red. Genes which targeting sgRNAs were enriched in population L (lowest ChFP/GFP ratio) are shown in blue. (**C**) TOP screen hits for each staller (red – KOs causing readthrough, blue – KOs increasing stalling). Genes in bold indicate genes that appeared as TOP screen hits for both stallers.

The poly-K screen yielded 96 gene knockouts enriched in population H (of these 19 were enriched more than 2-fold), and 56 gene knockouts enriched in population L (and of these 26 were more than 2-fold) (Figure 4B, Table S3). The 102×PR screen yielded 34 gene knockouts enriched in population H (and of these 5 were more than 2-fold) and 43 gene knockouts enriched in population L (and of these 15 were more than 2-fold). 21 gene knockouts were observed in common for both screens that increased apparent read-through (H) and there were 12 genes in common that appeared to increase stalling (L). Figure 4C shows a further refined list of top gene knockouts for each staller which consists of gene knockouts that were significantly enriched in one population (FDR<0.05, LFC>0) and simultaneously depleted from the opposite population (FDR<0.05, LFC<0).

The genes found in the poly-K screen included *ZNF598*, a previously known activator of the RQC (36), and members of ASC-1 complex (*ASCC3*, *ASCC2* and *TRIP4*) that disassembles collided ribosomes that result from stalling on polyadenosine mRNA (49) (Table 1). These knockouts were enriched in population H, which is consistent with other studies showing their knockout to improve poly-K readthrough (36,37,49). Another gene involved in RQC activation, *RACK1* (37), also appeared in our screen. *RACK1* knockouts were significantly depleted from population L and on the threshold for significant enrichment in population H (a false discovery rate value of 0.059). Collectively, these findings provided confidence in the screen for finding the relevant knockouts that regulate mechanisms involved in sensing and resolving stalled ribosomes. It was previously shown that EIF4E2-GIGYF2 (also known as 4EHP-GIGYF2) complex represses translation on defective mRNAs (50) and is recruited to ribosome collisions along with ZNF598 (51). According to our screen, cells with *EIF4E2* and *GIGYF2* knockouts appeared to have more stalling with the poly-K staller. This finding supports the hypothesis that in the absence of EIF4E2-GIGYF2 complex ZNF598 function on polyadenosine mRNA is enhanced.

**Table 1.**
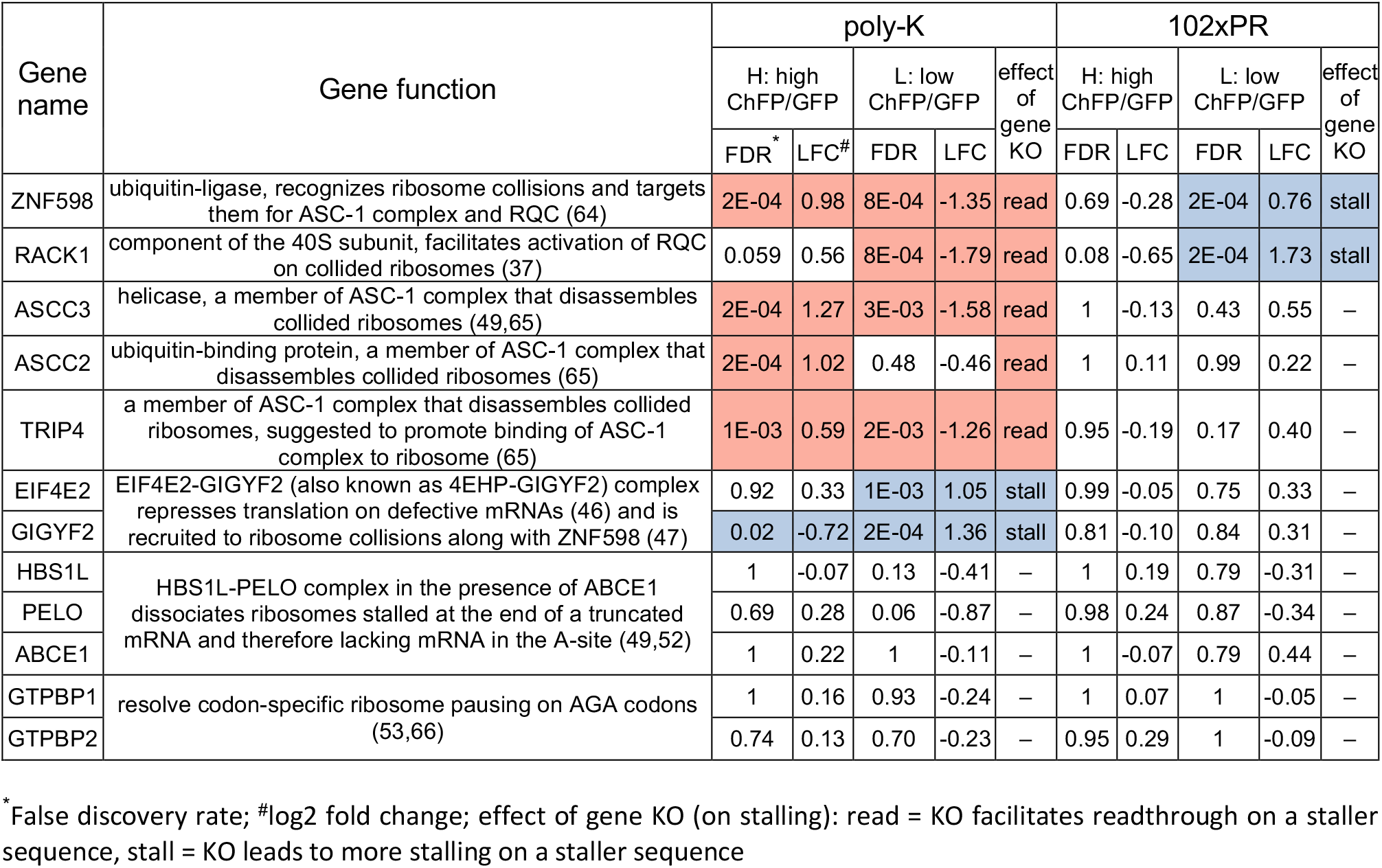
Enrichment factors for genes involved in rescue of stalled ribosomes.

| Gene name | Gene function | poly-K |  |  |  |  | 102xPR |  |  |  |  |
| --- | --- | --- | --- | --- | --- | --- | --- | --- | --- | --- | --- |
|  |  | H: high ChFP/GFP |  | L: low ChFP/GFP |  | effect of gene KO | H: high ChFP/GFP |  | L: low ChFP/GFP |  | effect of gene KO |
|  |  | FDR* | LFC# | FDR | LFC |  | FDR | LFC | FDR | LFC |  |
| ZNF598 | ubiquitin-ligase, recognizes ribosome collisions and targets them for ASC-1 complex and RQC (64) | 2E-04 | 0.98 | 8E-04 | -1.35 | read | 0.69 | -0.28 | 2E-04 | 0.76 | stall |
| RACK1 | component of the 40S subunit, facilitates activation of RQC on collided ribosomes (37) | 0.059 | 0.56 | 8E-04 | -1.79 | read | 0.08 | -0.65 | 2E-04 | 1.73 | stall |
| ASCC3 | helicase, a member of ASC-1 complex that disassembles collided ribosomes (49,65) | 2E-04 | 1.27 | 3E-03 | -1.58 | read | 1 | -0.13 | 0.43 | 0.55 | – |
| ASCC2 | ubiquitin-binding protein, a member of ASC-1 complex that disassembles collided ribosomes (65) | 2E-04 | 1.02 | 0.48 | -0.46 | read | 1 | 0.11 | 0.99 | 0.22 | – |
| TRIP4 | a member of ASC-1 complex that disassembles collided ribosomes, suggested to promote binding of ASC-1 complex to ribosome (65) | 1E-03 | 0.59 | 2E-03 | -1.26 | read | 0.95 | -0.19 | 0.17 | 0.40 | – |
| EIF4E2 | EIF4E2-GIGYF2 (also known as 4EHP-GIGYF2) complex represses translation on defective mRNAs (46) and is recruited to ribosome collisions along with ZNF598 (47) | 0.92 | 0.33 | 1E-03 | 1.05 | stall | 0.99 | -0.05 | 0.75 | 0.33 | – |
| GIGYF2 |  | 0.02 | -0.72 | 2E-04 | 1.36 | stall | 0.81 | -0.10 | 0.84 | 0.31 | – |
| HBS1L | HBS1L-PELO complex in the presence of ABCE1 dissociates ribosomes stalled at the end of a truncated mRNA and therefore lacking mRNA in the A-site (49,52) | 1 | -0.07 | 0.13 | -0.41 | – | 1 | 0.19 | 0.79 | -0.31 | – |
| PELO |  | 0.69 | 0.28 | 0.06 | -0.87 | – | 0.98 | 0.24 | 0.87 | -0.34 | – |
| ABCE1 |  | 1 | 0.22 | 1 | -0.11 | – | 1 | -0.07 | 0.79 | 0.44 | – |
| GTPBP1 | resolve codon-specific ribosome pausing on AGA codons (53,66) | 1 | 0.16 | 0.93 | -0.24 | – | 1 | 0.07 | 1 | -0.05 | – |
| GTPBP2 |  | 0.74 | 0.13 | 0.70 | -0.23 | – | 0.95 | 0.29 | 1 | -0.09 | – |

\* False discovery rate; #log2 fold change; effect of gene KO (on stalling): read = KO facilitates readthrough on a staller sequence, stall = KO leads to more stalling on a staller sequence

Genes known to be involved in activating the RQC showed a distinct enrichment pattern in the 102×PR screen (Table 1). Notably, *ZNF598* and *RACK1* knockouts were enriched in population L which is the opposite to what we observed for poly-K staller. *ASCC3*, *ASCC2*, *TRIP4* knockouts showed no enrichment in the 102×PR screen despite being hits in the poly-K screen. Other ribosome rescue factors seen in the polyK screen that can dissociate elongation complexes stalled at the 3` end of mRNA (*HBS1L*, *PELOTA*, *ABCE1*) (52), interact with ribosome collisions (*EIF4E2*, *GIGYF2*) or rescue ribosomes stalled at AGA arginine codons (*GTPBP1*, *GTPBP2*) (53) were not enriched in the 102×PR screen. These results suggested that co-translation and ribosome-associated quality control mechanisms that recognize and eliminate arrested ribosomes on polyadenosine sequences or other defective mRNAs might be unable to perform this function for poly-PR-induced stalls.

For further clues to pathways associated with genes identified in our screen we performed functional enrichment analysis. For both stallers, GO terms related to translation were enriched among gene knockouts improving the readthrough (Figure 5A–B). These are consistent with the key role of global translation since interference with translation strongly impacted the stall reporter behavior in our screen. This finding is supported by prior work in yeast showing that deletion of ribosomal proteins increases readthrough of poly-K and no-go decay (NGD) reporters (13). Knockouts of elongator complex (*ELP3*, *ELP4*, *ELP5*) and thiourydilase *CTU1,* which are required to efficiently decode AAA repeats by modifying U34 of lysyl-tRNA(UUU) (54), decreased the readthrough of poly-K. In addition, deletion of lysyl-tRNA synthetase (*KARS*) which catalyzes the formation of Lys-tRNA^Lys^ selectively increased stalling on poly-K, but not on 102×PR staller, which lacks Lys codons. The identification of these genes and pathways provide evidence the screen can identify proteins involved in both improving and reducing readthrough. Of note, genes activating RQC were identified in both screens, yet the genes showed different enrichment patterns further supporting our conclusions thus far that the surveillance and quality control mechanisms of stalling is dissimilar between polyadenosine and mRNA encoding 102×PR. Lastly, the enrichment of nuclear transport and nuclear envelope organization genes in 102×PR screen may relate to the recruitment of the poly-PR to the nucleus.

**Figure 5.**
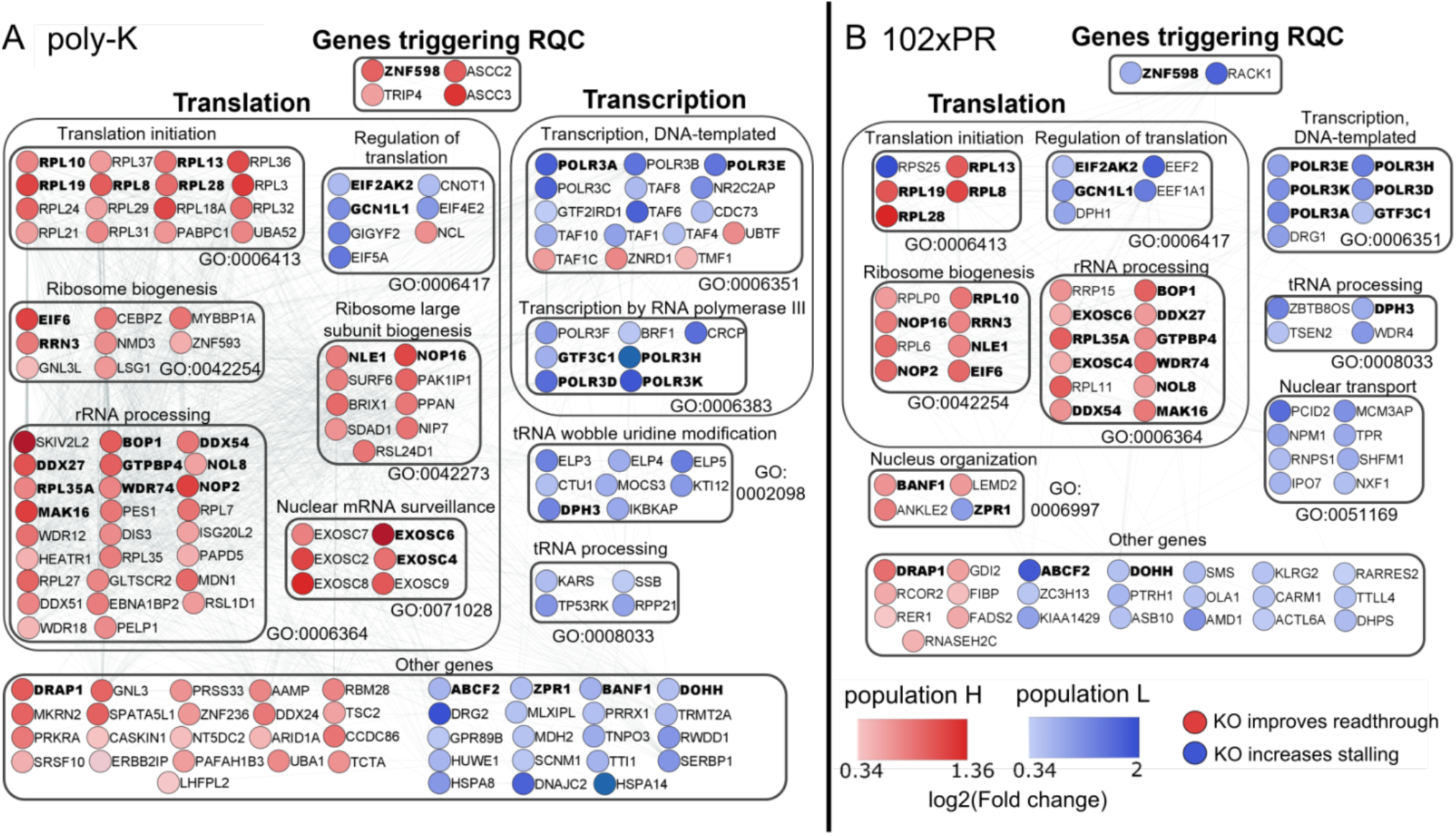
Top enriched gene ontology (GO) terms for screen hits at 5% FDR. (**A**) poly-K, (**B**) 102×PR. Genes in bold were the screen hits for both stallers.

Next, we further examined the role of 14 genes identified in our screen by creating individual knockout cell pools (Figure 6, Table 2). These genes were chosen on the basis of having large log2 fold change values in the knockout screen (*DRG2* and *SKIV2L2* for poly-K; *NPM1* and *RPS25* for 102×PR; *ABCF2*, *BOP1*, *DDX27*, *EIF2AK2*, *EIF6*, *NOP2* and *RPL19* for both stallers), or having opposing effects between polyK and poly-PR (*ZNF598*, *RACK1* and *BANF1*). To understand off-target impacts on the stall reporter readout, we included a control “linker” construct that contained a sequence known to not induce stalling (35–38). ZNF598 and RACK1 have been previously shown to increase readthrough of poly-K coded by (AAA) codons when their levels are reduced (36,37). Accordingly, their knockout increased the ChFP to GFP ratio by 3.7- and 2.7-fold, respectively, for the poly-K staller. By contrast, their knockout decreased the ratio for the 102×PR staller, in agreement with the screen outcomes which indicated that stalling was more pronounced when these genes are depleted. All other genes knockouts that impacted ChFP/GFP ratio on poly-K staller mostly had negligible or small effects on the corresponding ratios for the 102×PR readthrough. Collectively, these results indicated that the surveillance mechanisms involved in sensing stalls on polyadenosine mRNA have minimal effectivity in managing the stalls conferred by the 102×PR construct even if they may be able to sense the stalls as problematic based on their upregulation in the RNA-seq data.

**Figure 6.**
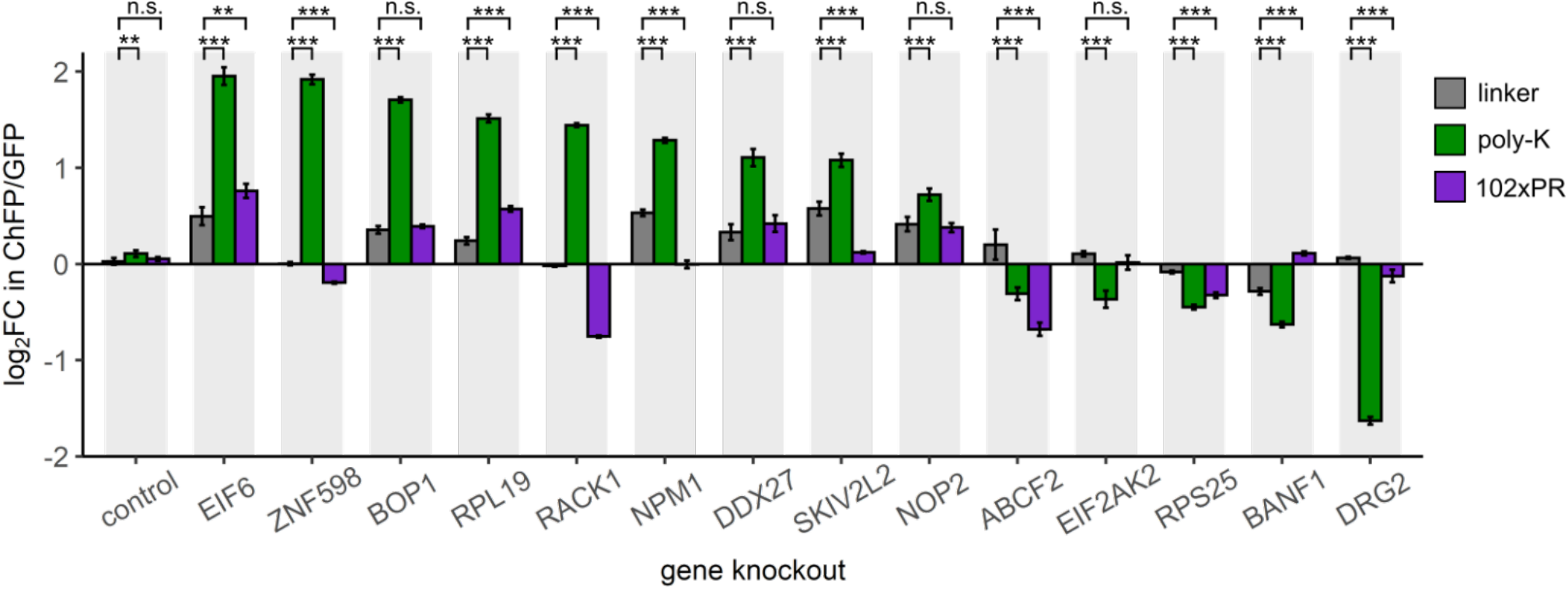
Log2 fold changes in ChFP/GFP ratio after KO of different genes selected from our screen hits. Control – cells expressing non-targeting sgRNA. Bars show means and SD, n=4. For each gene knockout, log2 fold changes for both stallers were compared to log2 fold changes for a linker construct using one-way ANOVA and Dunnett’s post hoc test. n.s. – not significant; ** P<0.01; *** P<0.0001.

**Table 2.** Validation of screen hits by individual knockouts.

| gene | function | KO effect on staller readthrough |  |  |  |  |  |
| --- | --- | --- | --- | --- | --- | --- | --- |
|  |  | in screen<sup>1</sup> |  | individual KO cell pool, compared to control cells<sup>2</sup> |  | individual KO cell pool, compared to linker<sup>3</sup> |  |
|  |  | poly-K | 102xPR | poly-K | 102xPR | poly-K | 102xPR |
| ABCF2 | unknown, a member of ABC transporters family | stall | stall | stall *** | stall *** | stall *** | stall *** |
| BANF1 | nuclear assembly and chromatin organization (67) | stall | read | stall *** | n.s. | stall *** | read *** |
| BOP1 | 60S ribosome subunit biogenesis (68) | read | read | read *** | read *** | read *** | n.s. |
| DDX27 | RNA helicase involved in rRNA processing (69) | read | read | read *** | read *** | read *** | n.s. |
| DRG2 | unknown, important role in cell growth (70) | stall | n.s. | stall *** | stall *** | stall *** | stall *** |
| EIF2AK2 | (also: PKR) inhibits cellular translation during stress (71) | stall | stall | stall *** | n.s. | stall *** | n.s. |
| EIF6 | ribosome biogenesis, binds to 60S to prevent premature association with 40S subunit (72) | read | read | read *** | read *** | read *** | read ** |
| NOP2 | nucleolar maturation and ribosome biogenesis (73) | read | read | read *** | read *** | read *** | n.s. |
| NPM1 | RNA transport and ribosome biogenesis (74) | n.s. | stall | read *** | n.s. | read *** | stall *** |
| RACK1 | scaffolding protein, component of 40S ribosome; activator of RQC on polyadenosine mRNA (37) | n.s. | stall | read *** | stall *** | read *** | stall *** |
| RPL19 | 60S ribosomal protein | read | read | read *** | read *** | read *** | read *** |
| RPS25 | 40S ribosomal protein, required for efficient RAN translation of C9orf72 repeats (75) | n.s. | stall | stall *** | stall *** | stall *** | stall *** |
| SKIV2L2 | (also: MTREX) RNA helicase, associated with the RNA exosome complex (76) | read | n.s. | read *** | n.s. | read *** | stall *** |
| ZNF598 | activator of RQC on polyadenosine mRNA (36) | read | stall | read *** | stall *** | read *** | stall *** |

<sup>1</sup> based on the screen results, read = significantly enriched in H population, stall = significantly enriched in L population.
<sup>2,3</sup> comparisons were made between log2fold changes (LFC) in ChFP/GFP. For each knockout, LFC = log2(the ratio in cells with sgRNA targeting an indicated gene divided by the ratio in non-infected cells without sgRNA). n.s. – not significant; \*\* P<0.01; \*\*\* P<0.0001, as determined by one-way ANOVA and Dunnett's post hoc test (n=4).
<sup>2</sup> based on the comparison between LFC for KO cell pool and control cell pool (cells with non-targeting sgRNA).
<sup>3</sup> based on the comparison between LFC for KO cell pool expressing an indicated staller and LFC for the same KO cell pool expressing linker sequence that does not cause stalling.

To examine whether these effects of gene targeting were specific to arginine in the DPR, we tested the 102×GR, which also causes stalling (17). None of the gene knockouts had any major effects on the ChFP/GFP ratio, suggesting that the stalling behaves similarly to that of 102×PR (Figure S3). However, the two knockouts that worsened stalling for 102×PR (*RACK1* or *ABCF2*) appeared to have no effect on 102×GR. This may be explained by the finding that 102×GR is more substantially stalled than 102×PR and hence has no capability to be further stalled (17) (Figure 1B, C).

### Inhibiting translation increases the readthrough of poly-K, but has limited or negative effects on R-rich DPRs translation

While the effect of the 14 gene knockouts suggested that the canonical regulatory mechanisms overseeing stalls on polyadenosine mRNA do not seem to effectively resolve the R-rich DPR stalls, we did note that *EIF6* and *RPL19* knockouts yielded slightly increased ChFP/GFP ratios for both poly-K and poly-PR relative to the linker control (Figure 6). eIF6 is a translation initiation factor that binds 60S ribosomal subunits which prevents them premature joining with 40S subunits. Reduction of eIF6 levels causes a decrease in the amount of available 60S subunits and impairs protein synthesis (55). We therefore hypothesized that eIF6 knockout reduces ribosome load on mRNA which, according to the collision-based model, should mitigate arrest on stall sequences.

To test whether reducing the ribosome load and translational throughput on mRNA can allow underlying attenuated mechanisms capacity to correct the stalling phenotype, we performed a study modeled on prior experiments showing that reducing the number of ribosomal collisions increases protein output from some stall sequences and stabilizes their translation (13). First, we examined low concentrations of translation initiation inhibitor harringtonine to reduce the density of ribosomes on mRNAs (56). Harringtonine treatment led to an increase in ChFP/GFP ratios for poly-K, suggesting it increases its readthrough by reducing the occurrences of ribosome collisions (Figure 7A). This treatment also increased ChFP/GFP ratios for 102×PR, albeit to a lower extent, suggesting that while slowing ribosome density can be helpful to reduce 102×PR stalling, mechanisms to address the stalls appear attenuated if they are indeed present. To explore this phenomenon with different approaches, we partly inhibited translation elongation with cycloheximide which appeared to improve the readthrough of the poly-K, however it led to a reduced apparent readthrough for the 102×PR (Figure 7B). Investigation of these treatments with other R-rich DPRs showed that the effects of reducing stalling were restricted to the longest repeat lengths we tested (102×) and was more effective for PR compared to GR (Figure S4).

**Figure 7.**
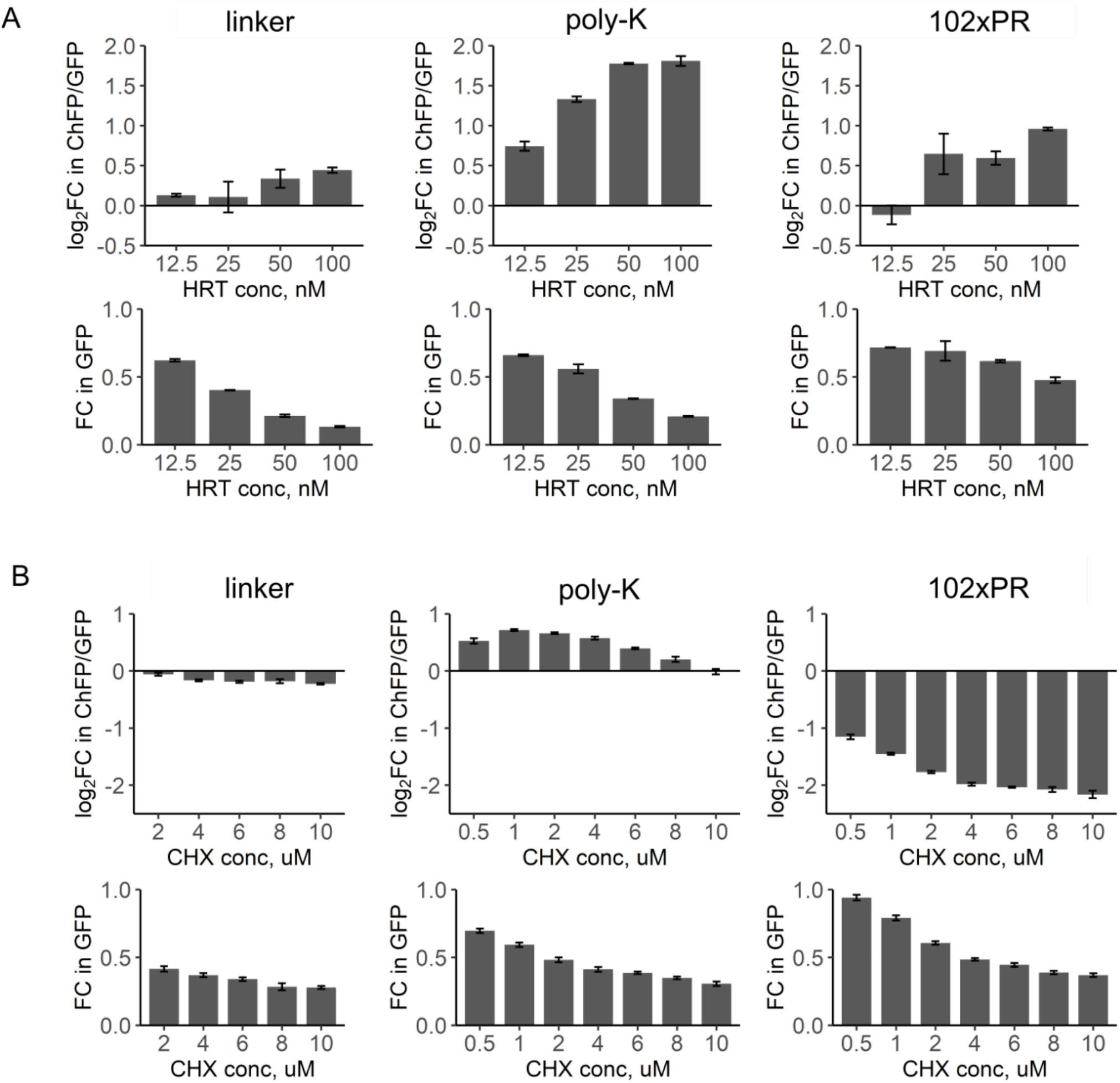
The effect of translation inhibition on the readthrough of different stallers. HEK293T cells were transfected with different fluorescent stall reporters and various concentrations of translation inhibitor were added 6 h after transfection before visible formation of fluorescent proteins. Cells were subjected to FACS after 18 h of treatment. Top plots on each panel show log2 fold change in ChFP/GFP ratio in cells treated with indicated concentration of translation inhibitor compared to untreated cells. Bottom plots on each panel show fold change in GFP fluorescence compared to untreated cells. Bars show means and SD, n=3. (**A**) Harringtonine, inhibitor of translation initiation. (**B**) Cycloheximide, inhibitor of translation elongation.

## DISCUSSION

Collectively, the findings presented here support a mechanism by which R-rich DPRs lead to stalling and are unable to be resolved by established quality control mechanisms that address polyadenosine-mediated stalling. This is in agreement with previous studies showing that RQC does not regulate the expression of endogenous proteins with polybasic sequences (57). The implications of these findings point to R-rich DPR-mediated stalling having no natural regulatory response, and therefore the stalled ribosomes may pose a particular danger to cellular homeostasis and viability. Notably, stalling on long poly-PR proteins is exacerbated when the global translation in compromised by adding elongation inhibitor. A recent study found that ageing, which is accompanied by a decline in cellular proteostasis, causes greater ribosome pausing at polybasic stretches (58). This suggests that a limited capacity that cells have to cope with stalls induced by a long poly-PR protein may be worse upon ageing and/or in the presence of other stresses, and therefore could prove as critical in the biological pathways leading to disease onset and development.

A question that remains unsolved is why the stalls occur in the first place. One possibility is that emergent nascent chains of the R-rich DPRs can interfere in trans with other ribosomes in addition to directly blocking the ribosome during synthesis. Poly-K can efficiently stall when coded with 21 consecutive AAA codons, which would render the sequence occupying maximally about two thirds of the ribosome exit tunnel (36). The ribosome exit tunnel can hold about 30 amino acids of a linear peptide chain and up to 60 amino acids for a peptide in an α-helical conformation (59). R-rich DPRs require more than 10 repeats (i.e. 20 amino acids for the poly-GR) and up to 40 repeats (80 amino acids for the poly-PR) to cause small levels of stalling (i.e. 25% or less). Such lengths would be sufficient to fill the ribosome exit tunnel and stall through an electrostatic jamming mechanism as previously proposed (17–19). However, the most effective stalling required longer dipeptide lengths (>30 repeats for poly-GR (60 amino acids) and >50 repeats for poly-PR (100 amino acids)) to stall more than 50%, and indeed did so in a length-dependent manner, suggesting that emergent DPR chain is involved in stalling. A preprint study suggested that the addition of short synthetic 20×PR and 20×GR polypeptides strongly inhibited global translation in vitro and occluded the exit tunnel of eukaryotic 80S ribosomes (19). Accordingly, it is reasonable to predict that during R-rich DPR translation emergent poly-GR or poly-PR nascent chains could bind and block the exit tunnels of trailing ribosomes on the mRNA leading to obstruction of translation in trans, and a collateral aggregation of translational machinery. The support to this mechanism comes from poly-GR and poly-PR interactomes being enriched with translational proteins (17,43). A further twist to this mechanism comes from simulations suggesting that the R-rich DPRs can phase separate once they reach a polymer length of about 25 repeats (60). Hence translation may be inhibited through a coalescence of ribosomes and blockage of their exit tunnels, which would presumably confer a great deal of toxic stress to cellular functioning.

Another aspect to our data is why the poly-GR appears to be more potent at stalling than poly-PR. One possibility is that the glycine allows greater conformational flexibility than proline (60). One consequence is that it may allow compact structures to accumulate in the ribosome exit tunnel, which may exacerbate blockage through electrostatic interactions. Indeed, the exit tunnel is wide enough to allow small protein domains to fold before exiting the ribosome tunnel (61). A greater flexibility may also lend to greater potency in influencing ribosomes in trans.

Poly-PR expression invokes a profound response from cells compared to poly-K expression, as reflected by substantial changes in mRNA levels for thousands of genes including genes involved in translation and cytoskeleton. Some of these responses likely arise from the multi-pronged toxicity previously implicated from poly-PR, including inhibition of translation (42–44) and the disruption of cytoskeleton architecture (17,45,62). Other responses relate more directly to stress response pathways including ribosome quality control (RQC), integrated stress response (ISR), unfolded protein response (UPR) and heat shock response activator HSF1. The activation of RQC pathway is important since it suggests that mechanisms exist to recognize the stalls, yet our other findings suggest that they are unable to be adequately resolved. Indeed other work has pointed to a role for RQC protein ZNF598 promoting the cleavage of poly-GR (but not poly-PR) by the ubiquitin-proteasome system while not affecting the readthrough efficiency (18). As such, the stalling may simply be overwhelming or improperly sensed by machinery that has not evolved to deal with such unnatural stalling sequences. In turn, it is plausible that the other stress responses are invoked in response to that inability to resolve through the RQC. Of note are studies finding DPR deposits in brain tissue to correlate with markers of the UPR (63).

## DATA AVAILABILITY

The data from RNA sequencing and genome-wide knockout screening is available in Tables S2 and S3 respectively. RNA sequencing data from this study have been deposited to NCBI GEO with the accession code: GSE193962. All data supporting the findings of this study is available from the corresponding author upon request.

## CONFLICT OF INTEREST STATEMENT

None declared.

## ACKNOWLEDGEMENTS

This work was funded by grants to DMH (National Health and Medical Research Council APP1161803 and Australian Research Council DP170103093) and to LF (Department of Health and Human Services acting through the Victorian Cancer Agency (fellowship MCRF16007)).

## Notes

### Competing Interest Statement

The authors have declared no competing interest.

