## Supplementary Figures for "Arginine-rich C9ORF72 ALS Proteins Stall Ribosomes in a Manner Distinct From a Canonical Ribosome-Associated Quality Control Substrate"

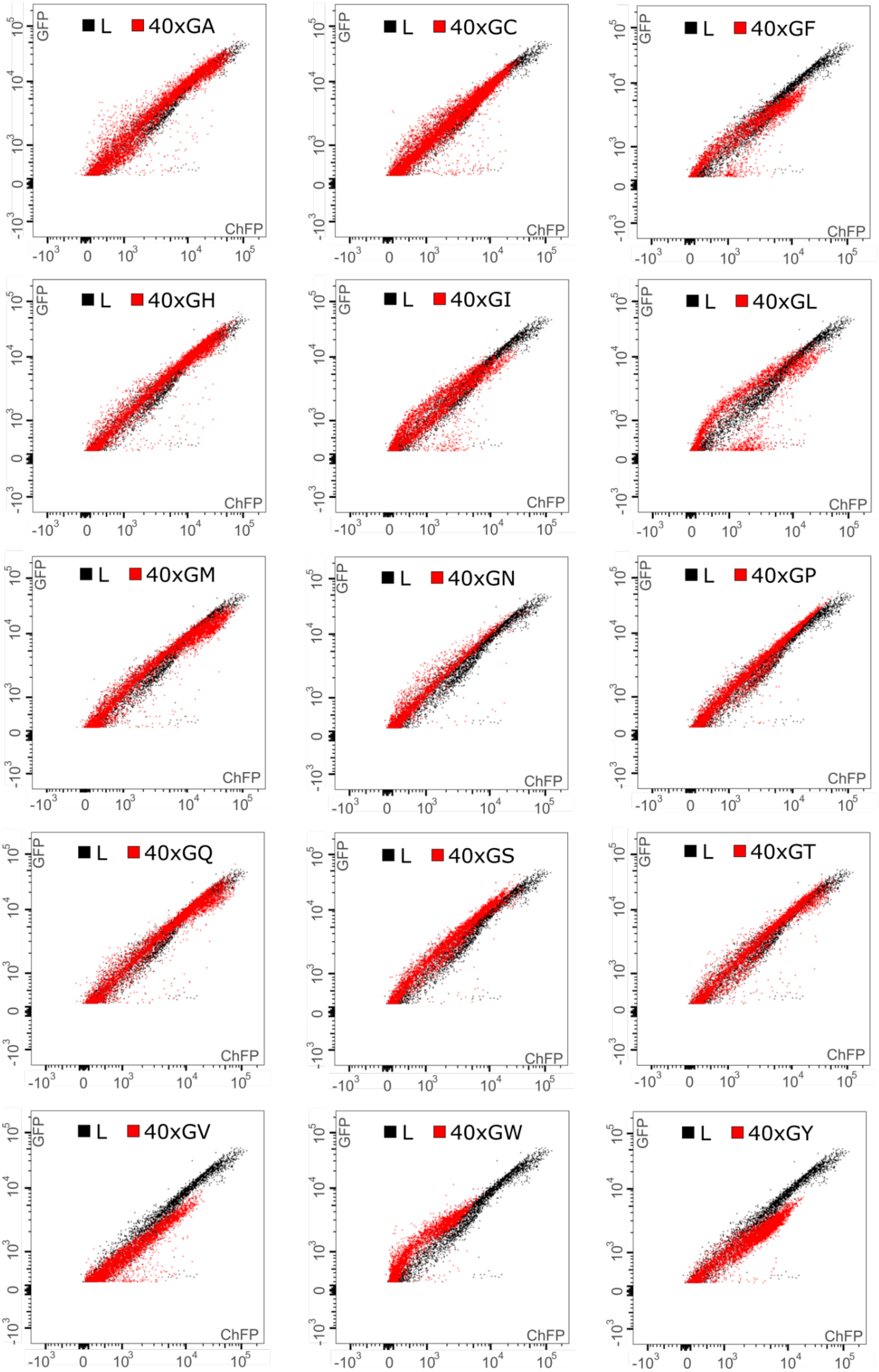
L
40xGA
40xGC
40xGF
40xGH
40xGI
40xGL
40xGM
40xGN
40xGP
40xGQ
40xGS
40xGT
40xGV
40xGW
40xGY
GFP
ChFP
$-10^3$
0
$10^3$
$10^4$
$10^5$

**Figure S1. FACS plots of cells expressing various DPRs.**. Fluorescent reporters containing indicated dipolypeptide repeat sequences (40×GX, where X – every canonical amino acid) between GFP and ChFP were transfected into 293T cells and analyzed by flow cytometry. The resulting cellular GFP and ChFP levels are depicted in the fluorescence-activated cell sorting (FACS) plots. The linker sequence (non-staller control) is shown in grey on each flow chart for the comparison.

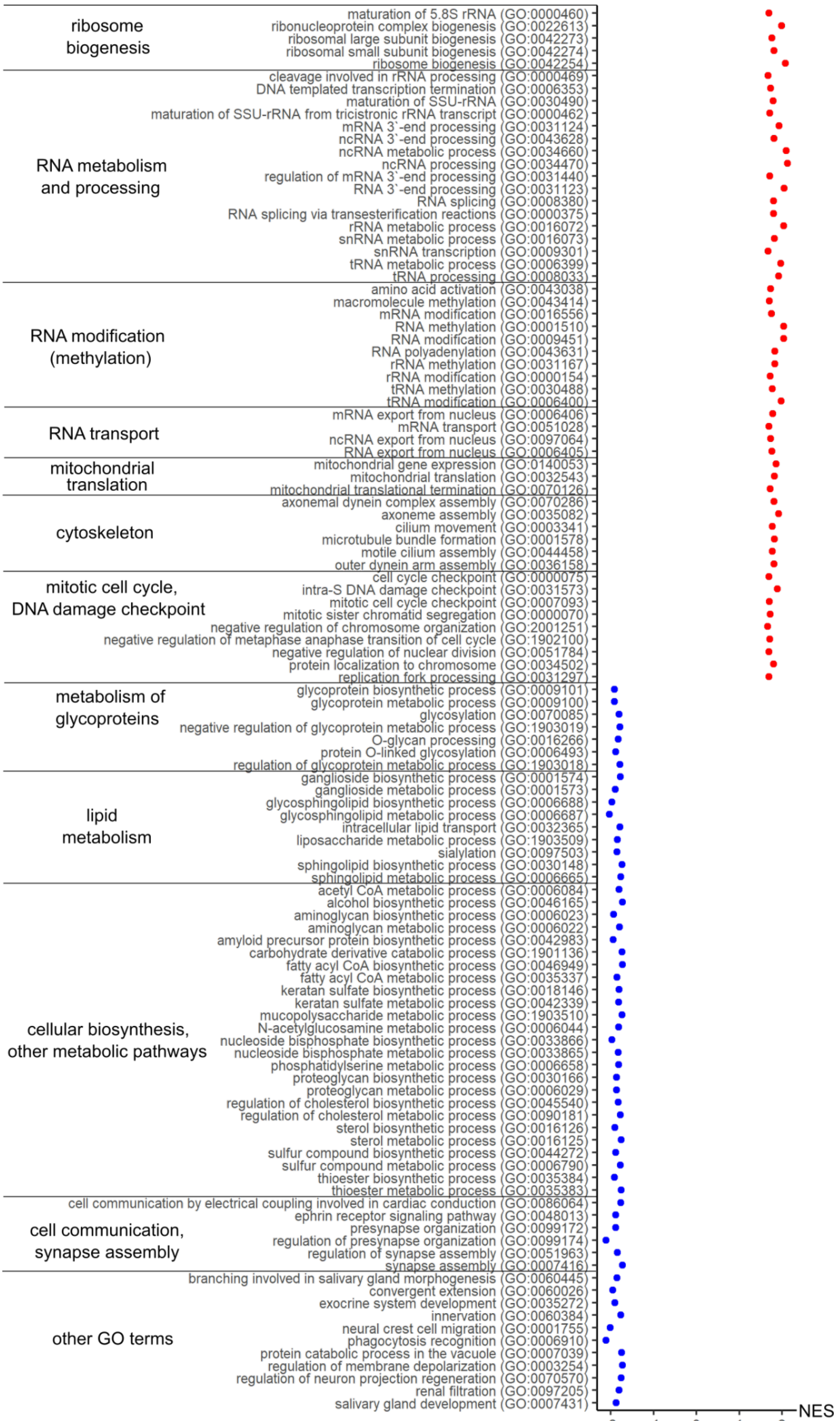

ribosome biogenesis
maturation of 5.8S rRNA (GO:0000460)
ribonucleoprotein complex biogenesis (GO:0022613)
ribosomal large subunit biogenesis (GO:0042273)
ribosomal small subunit biogenesis (GO:0042274)
ribosome biogenesis (GO:0042254)
RNA metabolism and processing
cleavage involved in rRNA processing (GO:0000469)
DNA templated transcription termination (GO:0006353)
maturation of SSU-rRNA (GO:0030490)
maturation of SSU-rRNA from tricistronic rRNA transcript (GO:0000462)
mRNA 3`-end processing (GO:0031124)
ncRNA 3`-end processing (GO:0043628)
ncRNA metabolic process (GO:0034660)
ncRNA processing (GO:0034470)
regulation of mRNA 3`-end processing (GO:0031440)
RNA 3`-end processing (GO:0031123)
RNA splicing (GO:0008380)
RNA splicing via transesterification reactions (GO:0000375)
rRNA metabolic process (GO:0016072)
snRNA metabolic process (GO:0016073)
snRNA transcription (GO:0009301)
tRNA metabolic process (GO:0006399)
tRNA processing (GO:0008033)
RNA modification (methylation)
amino acid activation (GO:0043038)
macromolecule methylation (GO:0043414)
mRNA modification (GO:0016556)
RNA methylation (GO:0001510)
RNA modification (GO:0009451)
RNA polyadenylation (GO:0043631)
rRNA methylation (GO:0031167)
rRNA modification (GO:0000154)
tRNA methylation (GO:0030488)
tRNA modification (GO:0006400)
RNA transport
mRNA export from nucleus (GO:0006406)
mRNA transport (GO:0051028)
ncRNA export from nucleus (GO:0097064)
RNA export from nucleus (GO:0006405)
mitochondrial translation
mitochondrial gene expression (GO:0140053)
mitochondrial translation (GO:0032543)
mitochondrial translational termination (GO:0070126)
cytoskeleton
axonemal dynein complex assembly (GO:0070286)
axoneme assembly (GO:0035082)
cilium movement (GO:0003341)
microtubule bundle formation (GO:0001578)
motile cilium assembly (GO:0044458)
outer dynein arm assembly (GO:0036158)
mitotic cell cycle, DNA damage checkpoint
cell cycle checkpoint (GO:0000075)
intra-S DNA damage checkpoint (GO:0031573)
mitotic cell cycle checkpoint (GO:0007093)
mitotic sister chromatid segregation (GO:0000070)
negative regulation of chromosome organization (GO:2001251)
negative regulation of metaphase anaphase transition of cell cycle (GO:1902100)
negative regulation of nuclear division (GO:0051784)
protein localization to chromosome (GO:0034502)
replication fork processing (GO:0031297)
metabolism of glycoproteins
glycoprotein biosynthetic process (GO:0009101)
glycoprotein metabolic process (GO:0009100)
glycosylation (GO:0070085)
negative regulation of glycoprotein metabolic process (GO:1903019)
O-glycan processing (GO:0016266)
protein O-linked glycosylation (GO:0006493)
regulation of glycoprotein metabolic process (GO:1903018)
lipid metabolism
ganglioside biosynthetic process (GO:0001574)
ganglioside metabolic process (GO:0001573)
glycosphingolipid biosynthetic process (GO:0006688)
glycosphingolipid metabolic process (GO:0006687)
intracellular lipid transport (GO:0032365)
liposaccharide metabolic process (GO:1903509)
sialylation (GO:0097503)
sphingolipid biosynthetic process (GO:0030148)
sphingolipid metabolic process (GO:0006665)
cellular biosynthesis, other metabolic pathways
acetyl CoA metabolic process (GO:0006084)
alcohol biosynthetic process (GO:0046165)
aminoglycan biosynthetic process (GO:0006023)
aminoglycan metabolic process (GO:0006022)
amyloid precursor protein biosynthetic process (GO:0042983)
carbohydrate derivative catabolic process (GO:1901136)
fatty acyl CoA biosynthetic process (GO:0046949)
fatty acyl CoA metabolic process (GO:0035337)
keratan sulfate biosynthetic process (GO:0018146)
keratan sulfate metabolic process (GO:0042339)
mucopolysaccharide metabolic process (GO:1903510)
N-acetylglucosamine metabolic process (GO:0006044)
nucleoside bisphosphate biosynthetic process (GO:0033866)
nucleoside bisphosphate metabolic process (GO:0033865)
phosphatidylserine metabolic process (GO:0006658)
proteoglycan biosynthetic process (GO:0030166)
proteoglycan metabolic process (GO:0006029)
regulation of cholesterol biosynthetic process (GO:0045540)
regulation of cholesterol metabolic process (GO:0090181)
sterol biosynthetic process (GO:0016126)
sterol metabolic process (GO:0016125)
sulfur compound biosynthetic process (GO:0044272)
sulfur compound metabolic process (GO:0006790)
thioester biosynthetic process (GO:0035384)
thioester metabolic process (GO:0035383)
cell communication, synapse assembly
cell communication by electrical coupling involved in cardiac conduction (GO:0086064)
ephrin receptor signaling pathway (GO:0048013)
presynapse organization (GO:0099172)
regulation of presynapse organization (GO:0099174)
regulation of synapse assembly (GO:0051963)
synapse assembly (GO:0007416)
other GO terms
branching involved in salivary gland morphogenesis (GO:0060445)
convergent extension (GO:0060026)
exocrine system development (GO:0035272)
innervation (GO:0060384)
neural crest cell migration (GO:0001755)
phagocytosis recognition (GO:0006910)
protein catabolic process in the vacuole (GO:0007039)
regulation of membrane depolarization (GO:0003254)
regulation of neuron projection regeneration (GO:0070570)
renal filtration (GO:0097205)
salivary gland development (GO:0007431)
-2
-1
0
1
2
NES

**Figure S2. Significantly upregulated (red) and downregulated (blue) GO terms in 102xPR-expressing cells grouped into functional clusters.** NES – normalized enrichment score.

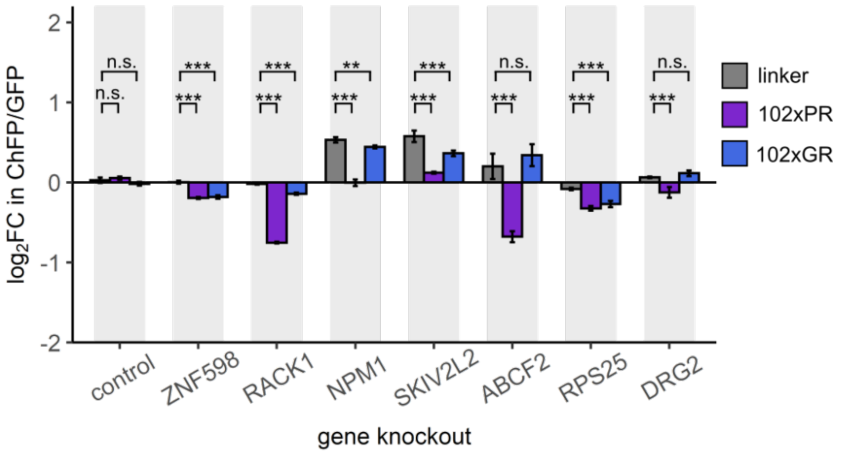


**Figure S3. Log$_2$ fold changes (LFC) in ChFP/GFP ratio for 102×PR and 102×GR stallers after knockout of different genes selected from our screen hits.** Control – cells expressing non-targeting sgRNA. Error bars show SD, n=4. For each gene knockout, log2 fold changes for both stallers were compared to log2 fold changes for a linker construct using one-way ANOVA and Dunnett's post hoc test. n.s. – not significant; ** $P<0.01$; *** $P<0.0001$.

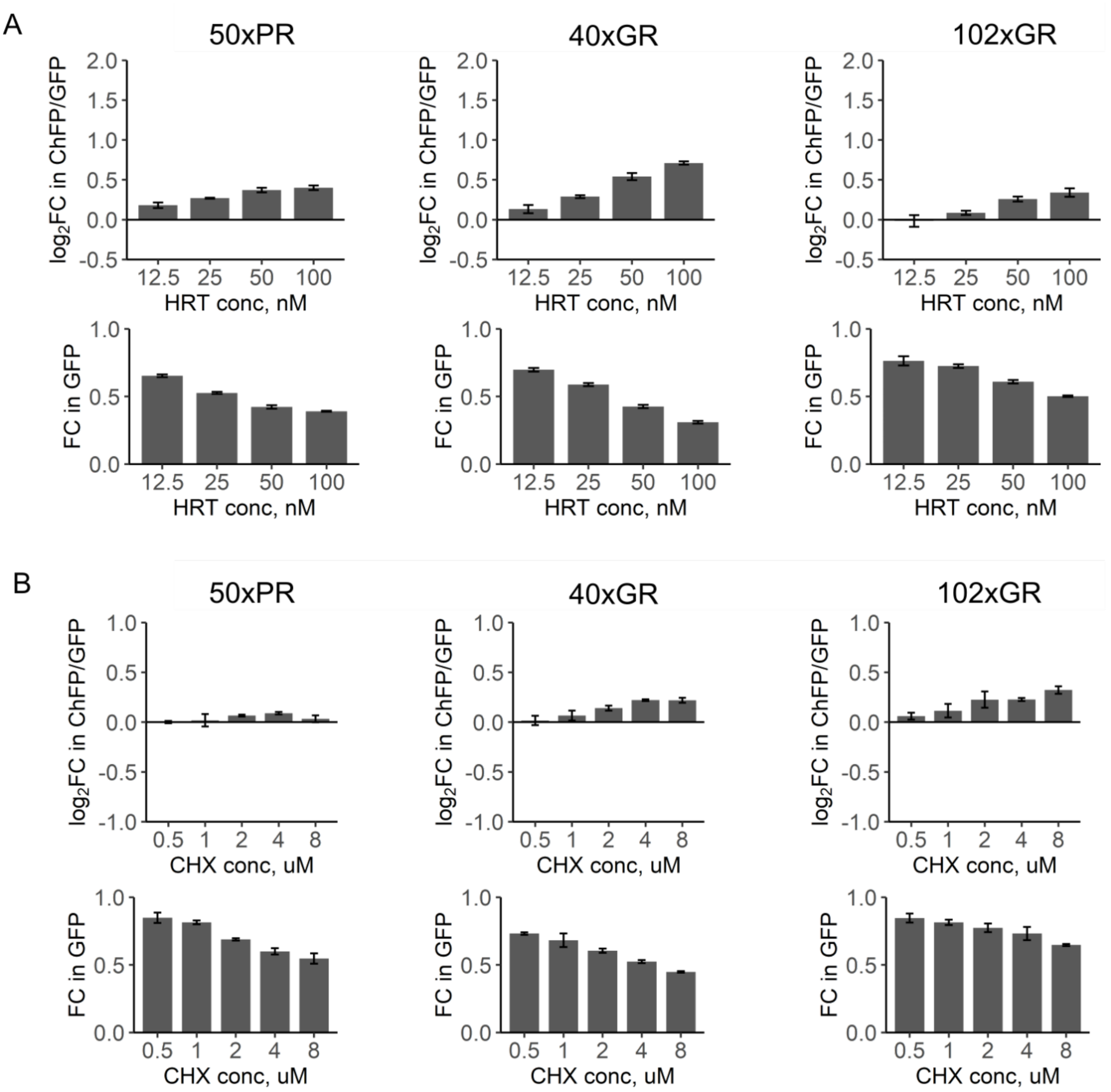


**Figure S4. The effect of translation inhibition on the readthrough of 50×PR, 40×GR and 102×GR.** HEK293T cells were transfected with different fluorescent stall reporters and various concentrations of translation inhibitor were added 6 h after transfection before visible formation of fluorescent proteins. Cells were subjected to FACS after 18 h of treatment. Top plots on each panel show log2 fold change in ChFP/GFP ratio in cells treated with indicated concentration of translation inhibitor compared to untreated cells. Bottom plots on each panel show fold change in GFP fluorescence compared to untreated cells. Error bars show SD, n=3. (**A**) Harringtonine, inhibitor of translation initiation. (**B**) Cycloheximide, inhibitor of translation elongation.
